# Engineered Terpenoid Production in *Synechocystis* sp. PCC 6803 Under Different Growth Conditions

**DOI:** 10.1101/2020.08.19.256149

**Authors:** Ryan A. Herold, Samantha J. Bryan

## Abstract

Terpenoids are the largest class of natural products and have applications in a wide variety of industries. Many terpenoids can be chemically synthesized or extracted from plants, but this is often uneconomical or unsustainable. An alternative production method relies on the heterologous expression of terpene synthase enzymes in cyanobacteria, producing the desired compounds directly from carbon dioxide. In this work, a patchoulol synthase enzyme from *Pogostemon cablin* (patchouli) was expressed in the cyanobacterium *Synechocystis* sp. PCC 6803 under four different growth conditions. Final yields of patchoulol from each growth condition were as follows: 249 μg L^−1^, photoautotrophic growth; 6.5 μg L^−1^, mixotrophic growth; 27.6 μg L^−1^, bicarbonate low light; 31.7 μg L^−1^, bicarbonate high light. By comparing patchoulol production across growth conditions, we identified a significant positive correlation between the production of photopigments (chlorophyll and carotenoids) and the production of patchoulol. Importantly, this relationship was found to be stronger than the correlation between cell density and patchoulol production across growth conditions, which was not statistically significant. The relationship between photopigments and patchoulol should be generalizable to the production of other terpenoids that rely on expression of the endogenous methylerythritol phosphate (MEP) pathway in cyanobacteria. Based on the results of this work, we propose a strategy for maximizing terpenoid production in cyanobacteria by optimizing growth conditions for photopigment production, resulting in increased flux through the MEP pathway. This strategy has the advantage of facile photopigment quantification using simple spectroscopic methods, and optimized growth conditions can be utilized in partnership with conventional terpenoid production strategies to further improve yields.

---

Cyanobacteria are a diverse group of photosynthetic prokaryotes that produce a wide variety of useful products and are capable of growing on carbon dioxide as a sole carbon source. Some cyanobacteria species are capable of fixing atmospheric nitrogen in addition to fixing carbon dioxide—an impressive feat that eliminates the need to add nitrogen-based fertilizers which require large amounts of energy to produce. In addition, many cyanobacteria can be grown in fresh or saltwater and do not compete with crop cultivation by requiring arable land, making them attractive organisms for the production of renewable fuels and chemicals.^1,2^ Although genetic tools for cyanobacteria are less developed than those for model heterotrophs such as yeast and *E. coli*, significant progress has been made toward characterizing genetic parts (e.g., promoters, ribosome binding sites, neutral sites, and recombineering plasmids) that make cyanobacteria more attractive chassis for industrial biotechnology.^3–8^ Much of that work has been done using *Synechocystis* sp. PCC 6803, hereinafter referred to as *Synechocystis*. *Synechocystis* is a model organism that has been used for studying photosynthesis,^9^ plastid evolution,^10^ and circadian rhythms^11^ and was the first photoautotroph to have its genome fully sequenced.^12^

A biotechnology application that cyanobacteria are particularly suited for is the production of terpenoids. This is because cyanobacteria naturally produce large quantities of terpenoids in the form of carotenoids and the phytol tail of chlorophyll.^13^ Terpenoids (also called isoprenoids) are the largest class of natural products^14^ and include terpenes—unsaturated hydrocarbons that consist of one or more isoprene units linked together—and derivatives of terpenes.^15^ Altogether, there are at least 30,000 terpenes and over 50,000 natural terpenoids.^14^ This incredible diversity results from the abundance of terpene synthase enzymes that exist in nature, many of which produce multiple products.^16^ Cyanobacteria produce terpenoids via the methylerythritol phosphate (MEP) pathway,^17^ which utilizes pyruvate and glyceraldehyde-3-phosphate (G3P) as precursor molecules. Using cyanobacteria to produce terpenoids that are traditionally derived from plants takes advantage of the higher photosynthetic efficiency of cyanobacteria versus land plants,^18–21^ as well as a much faster growth rate (*Synechocystis* doubling time can be as short as 5.13 hours),^22^ allowing for substantially shorter time between harvests. The case for using cyanobacteria to produce traditionally plant-derived chemicals is further strengthened in regions of the world, such as southeast Asia, where rainforest is destroyed in order to obtain arable land for the production of commodity chemicals such as natural rubber (polyisoprene).^23,24^

Concerns about climate change and unsustainable chemical production methods have spurred interest in the production of terpenoids using cyanobacteria, and there are many review articles that have collated data from these studies.^13,25–28^ Although some studies have reported impressive terpenoid yields from cyanobacteria of up to 1.26 g L^−1^ (isoprene),^29^ the production of commodity chemicals is still generally uneconomical due to the low prices of chemicals derived from fossil fuels. In order for cyanobacterial terpenoid production to be economically viable, the target molecules must be sufficiently high-value compounds such that they can be produced at a lower price than the current method of production. Examples of such compounds include pharmaceuticals, nutraceuticals, natural pigments, and flavor and fragrance compounds.

In this study, our goal was to produce the terpenoid, patchoulol (also called patchouli alcohol), using *Synechocystis* and test how different growth conditions affected production. To achieve this, we expressed a codon-optimized, heterologous patchoulol synthase gene from the patchouli plant, *Pogostemon cablin*, and quantified the amount of patchoulol produced each day under four different growth conditions. Patchoulol is a sesquiterpenoid (C_15_), a group of molecules derived from farnesyl diphosphate (FPP) that have long been important in the fragrance and cosmetic industries.^30^ It has a strong woody scent and is the main component of patchouli essential oil, which is commercially produced by steam distilling dried leaves of the patchouli plant.^31,32^ Patchouli is mainly grown in south-east Asia (particularly Indonesia), and the essential oil is the tenth most important in the world by mass with a price ranging from $61–$150 per kg.^32^

Bioengineered production of patchoulol has gained significant interest in recent years, and in 2014 the Swiss company Firmenich launched Clearwood™, a biosynthetic patchouli-like fragrance containing patchoulol produced using engineered yeast.^32^ A review of the literature reveals studies utilizing a wide variety of different organisms for patchoulol production. The most common is *Saccharomyces cerevisiae*,^33–36^ with the highest yield reporting an impressive 59.2 mg L^−1^ and 466.8 mg L^−1^ patchoulol for batch and fed-batch fermentation conditions, respectively.^33^ Another notable study utilized the heterotrophic bacterium, *Corynebacterium glutamicum*, which was able to achieve a yield of 60 mg L^−1^ from a strain that had been engineered through genomic deletions and *ispA* overexpression.^37^

Studies of more relevance to this work are those using photosynthetic organisms for patchoulol production. Though they generally provide lower yields than heterotrophs, photoautotrophs offer more sustainable chemical production. Studies utilizing photoautotrophs for patchoulol production include the eukaryotic microalgae, *Chlamydomonas reinhardtii*,^38^ and a moss, *Physcomitrella patens*,^39^ which provided yields of 922 μg g^−1^ and 1.34 mg g^−1^ dry weight, respectively. In *Synechocystis*, expression of the patchoulol synthase gene under a strong copper inducible promoter yielded 17.3 mg L^−1^ patchoulol.^40^ That study utilized a highly optimized small-scale High-Density Cultivation (HDC) system with nutrient rich media, bicarbonate, and increasing light intensity. The impressive patchoulol yield achieved reflects the high cell densities that can be reached using HDC; however, when grown under typical batch conditions, the yields were similar to those achieved for other terpenoids.

This study focused on how growth and metabolism can unlock the constraints on terpenoid production in *Synechocystis*. Specifically, we investigated how patchoulol productivity was affected when cells used different metabolic growth strategies. This was achieved by growing our engineered patchoulol-producing *Synechocystis* strain under four different growth conditions (photoautotrophic, mixotrophic, bicarbonate low light, and bicarbonate high light), tracking daily patchoulol levels, and analyzing the relationship between patchoulol production and metabolites which utilize the MEP pathway.

## RESULTS AND DISCUSSION

### Generation of a Patchoulol-producing *Synechocystis* Strain

A glucose-tolerant wild-type strain of *Synechocystis* sp. PCC 6803 was engineered for patchoulol production. The patchoulol-producing strain of *Synechocystis* (PATS) was created by knocking out the native *psbA2* open reading frame and replacing it with a synthetic, codon optimized patchoulol synthase gene from *Pogostemon cablin*. Protein expression using the *psbA2* locus of *Synechocystis* is a commonly used technique that was first established for the production of carotenoids.^41^ Following transformation, antibiotic selection, and growth in liquid culture, isolated PATS cultures were screened using the primers 6803psbA2_F and 6803psbA2_R (**Supplemental Table 1**). The primers annealed outside the site of recombination in the *Synechocystis* genome, producing a smaller (1721 bp) amplicon for the wild type and a larger amplicon for the PATS strain (3480 bp) (**Figure 1**). By using external primers, we were able to confirm when cultures were fully or only partially segregated by the presence of one or two bands. Additional evidence that transformation had been successful and that a patchoulol-producing strain had been generated was provided by the presence of a strong, patchouli-like fragrance originating from PATS cultures that was not present for wild-type cultures.

**Figure 1.**
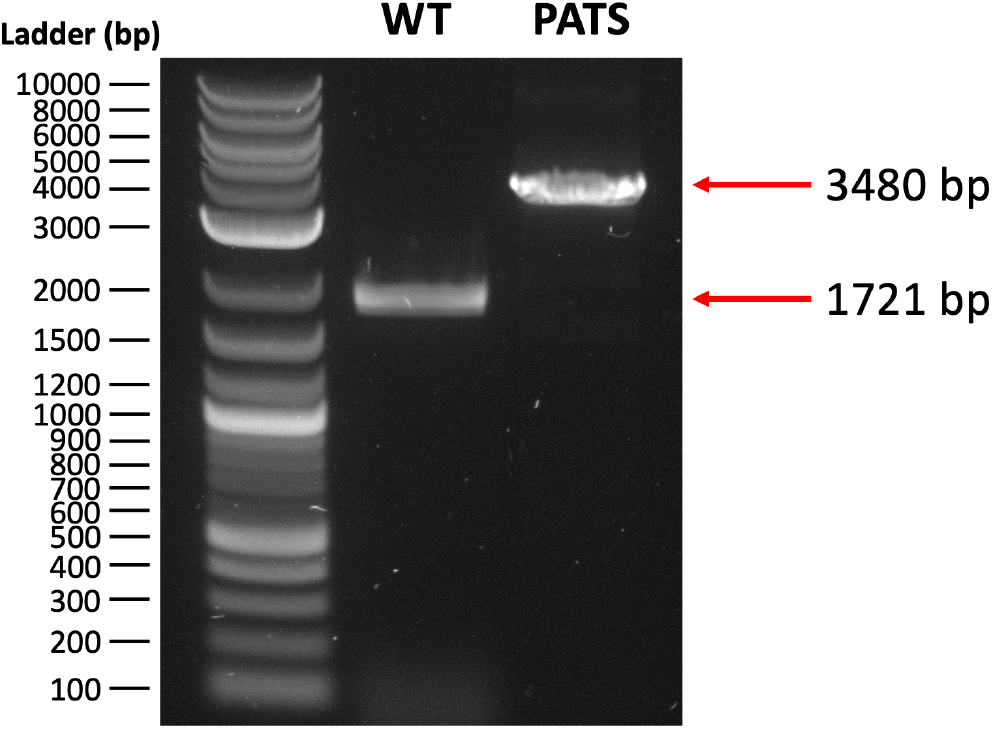
Agarose gel showing the fully segregated *Synechocystis* PATS strain compared to the wild type. DNA was amplified from cell boilate using primers that annealed in regions just outside the *psbA2* site where recombination occurred. PCR from the PATS and wild-type strains gave bands of the expected sizes, 3480 and 1721 bp, respectively.

To confirm that patchoulol was being produced by the isolated PATS strain, 70 mL cultures were grown photoautotrophically in a photobioreactor with a 10% (7 mL) n-dodecane (Acros Organics) solvent overlay to collect the excreted patchoulol, and the dodecane was analyzed via GCMS and compared to a patchoulol primary reference standard (**Figure 2**). The retention time of the patchoulol produced by *Synechocystis* matched perfectly with the patchoulol standard, and the mass spectra were identical and also matched the mass spectrum for patchoulol from the NIST Standard Reference Database.^42^

**Figure 2.**
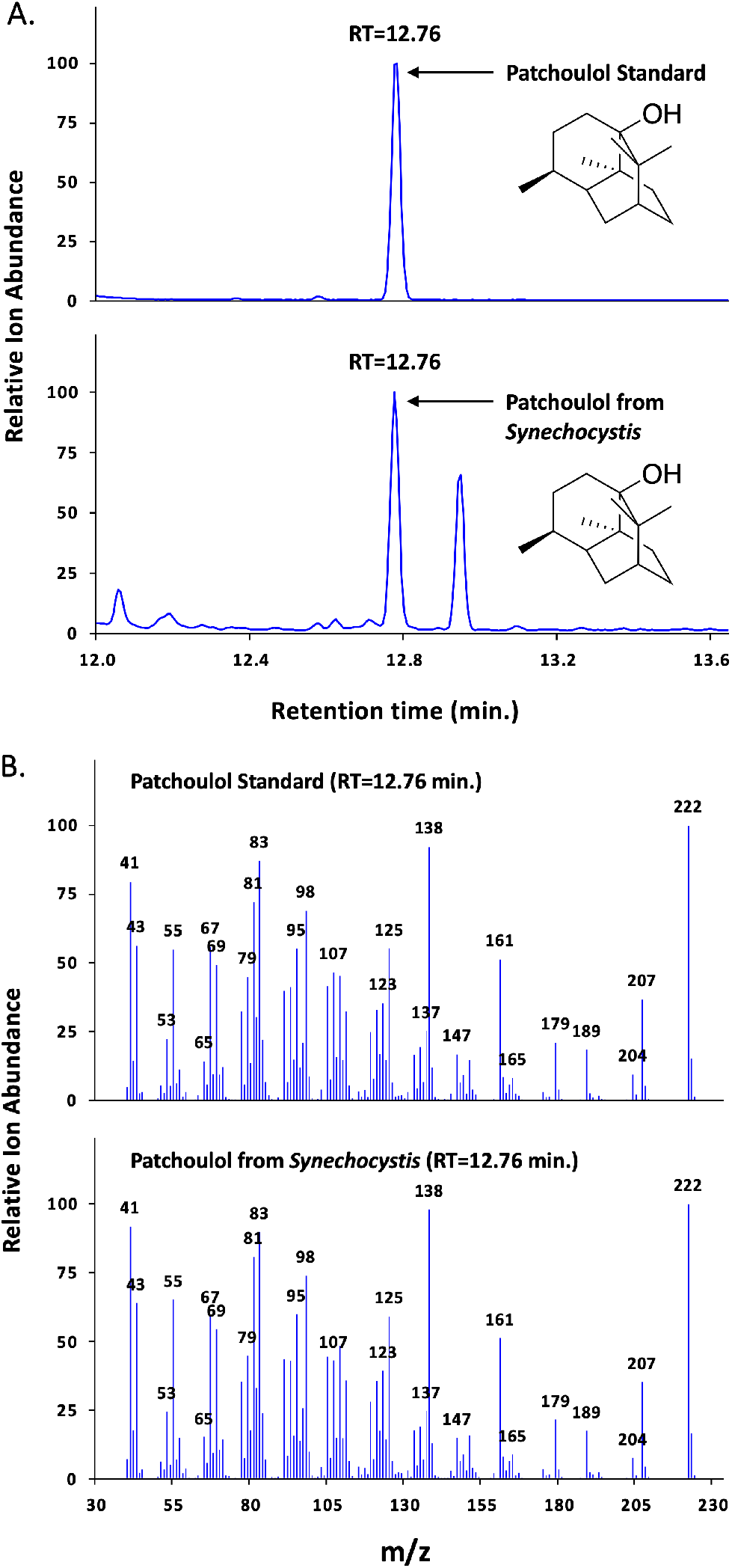
Gas total ion chromatograms and mass spectra of a patchoulol primary reference standard (Sigma-Aldrich) and patchoulol produced by *Synechocystis* in dodecane. (**A**) Gas chromatogram of patchoulol standard (top) and patchoulol produced by the recombinant *Synechocystis* PATS strain (bottom). (**B**) Mass spectra of patchoulol standard (top) and patchoulol produced by the *Synechocystis* PATS strain (bottom). Chromatograms and mass spectra were normalized to 100 to represent relative signal intensity.

### Photoautotrophic and Mixotrophic Patchoulol Production

Once we established that patchoulol was produced by the PATS strain, we tested how production differed when cultures were grown photoautotrophically versus mixotrophically. Photoautotrophic cultures were grown in a photobioreactor with 0.4% CO_2_ mixed with air as the sole carbon source. After 20 days of growth, the PATS strain produced an average final yield of 249 μg L^−1^ patchoulol (**Figure 3A)**, with a maximum specific production rate of 2.581 μg L^−1^ day^−1^ OD_750_^−1^ on day 14 (**Figure 3B)**. Specific production of patchoulol (daily production normalized to OD) was consistent after the first two days of growth, and there was no difference in growth between the wild-type and PATS strains (**Figure 3A**).

**Figure 3.**
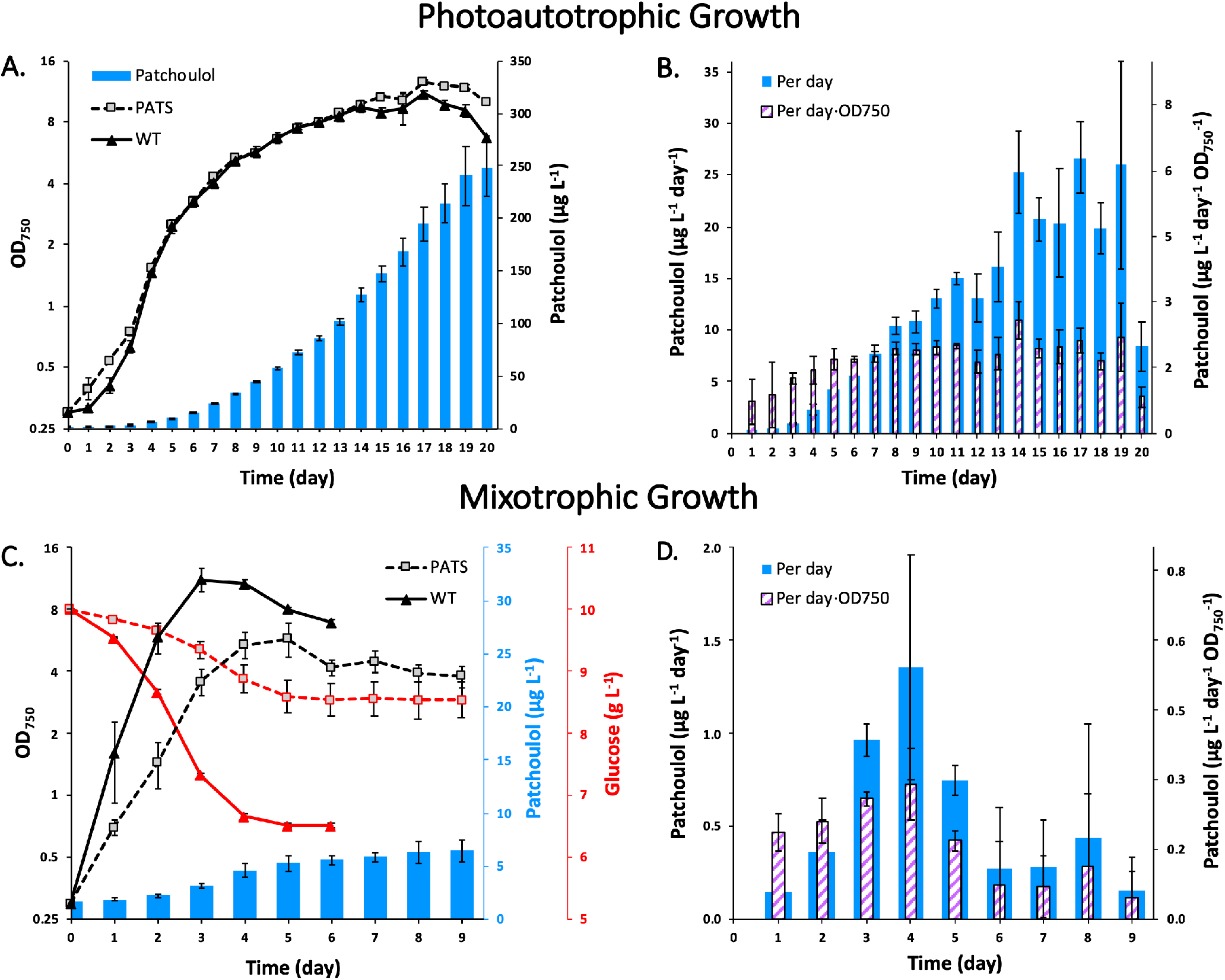
Photoautotrophic and mixotrophic production of patchoulol from *Synechocystis* cultures grown in a photobioreactor. (**A**) Growth curves of cultures grown photoautotrophically (0.4% CO_2_ mixed with air as sole carbon source) and patchoulol production by the PATS strain. (**B**) Daily productivity and specific productivity of PATS strain grown photoautotrophically. (**C**) Growth curves of cultures grown mixotrophically (0.4% CO_2_ + 10 g L^−1^ glucose) and patchoulol production by the PATS strain. Residual glucose was quantified at each sampling timepoint and is shown using a third y-axis (red). (**D**) Daily productivity and specific productivity of PATS strain grown mixotrophically. Error bars represent the standard deviation of triplicate cultures. **Note:** growth curves are plotted on a log base 2 scale to better represent exponential growth.

Cultures were grown under mixotrophic conditions with the hypothesis that patchoulol yields would increase relative to photoautotrophic conditions, as has been shown to be the case for the production of isobutanol.^18^ The same growth conditions were used for mixotrophic growth as for photoautotrophic growth, with the addition of 10 g L^−1^ glucose to the medium. Surprisingly, mixotrophic patchoulol production was much lower compared to photoautotrophic production. After nine days of growth, mixotrophic cultures produced an average of only 6.5 μg L^−1^ patchoulol (**Figure 3C)**, with a maximum specific production rate of 0.291 μg L^−1^ day^−1^ OD_750_^−1^ on day 4 (**Figure 3D**). Similar to patchoulol production under photoautotrophic conditions, specific productivity increased during the first few days of exponential growth. However, unlike photoautotrophic production, specific productivity of mixotrophic cultures declined immediately after peaking on day 4, and there was not an extended period of high patchoulol production that lasted multiple days. Additionally, mixotrophic conditions strongly inhibited growth of the PATS strain, which reached an average OD of only 4.53 compared to 11.17 for the wild type (**Figure 3C**). The reason for the growth inhibition is unknown; however, a preliminary experiment confirmed the growth inhibition for mixotrophic PATS cultures (**Supplemental Figure 1**). In most cases, the *psbA2* gene can be knocked out without deleterious effects,^4^ and it has been shown that the weakly expressed *psbA3* gene (which also codes for the D1 protein) is upregulated to compensate for the loss of *psbA2*.^43^ If the growth inhibition was caused by light-induced stress due to the lack of a functional *psbA2* gene, a similar growth inhibition should have also been observed under photoautotrophic conditions since the light intensity was the same.

In addition to tracking cell density and patchoulol production, residual glucose in the medium was quantified using HPLC (**Figure 3C**). The PATS strain used less than half as much glucose as the wild type, consistent with the cell densities observed. The wild-type and PATS strains used 3.49 g L^−1^ and 1.46 g L^−1^ of glucose, respectively. In order to better understand some of the metabolic differences between the strains under different growth conditions, chlorophyll and carotenoids were each quantified at two timepoints (early and late) during growth (**Figure 4**). Because both carotenoids and the phytol tail of chlorophyll are terpenoids produced from derivatives of FPP (the substrate for patchoulol synthase), their respective enzymes required for synthesis compete with patchoulol synthase for substrate. Therefore, we were also interested to see if the expression of patchoulol synthase reduced chlorophyll and/or carotenoid levels compared to the wild type.

**Figure 4.**
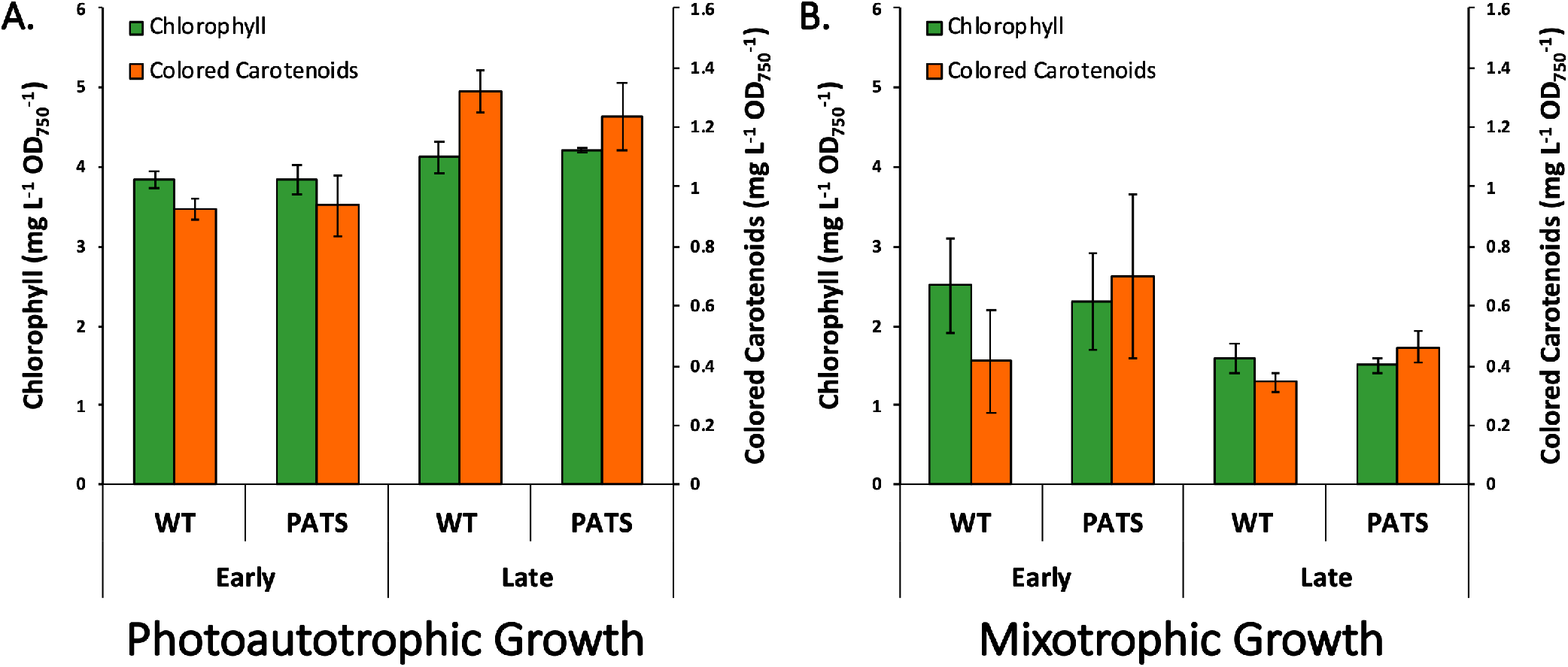
Chlorophyll and colored carotenoid analysis of wild-type and PATS *Synechocystis* strains grown in a photobioreactor. (**A**) Cultures grown photoautotrophically on 0.4% CO_2_. Samples for the early and late timepoints during growth were taken on days 4 and 13, respectively. (**B**) Cultures grown mixotrophically with 10 g L^−1^ glucose and 0.4% CO_2_. Early timepoint samples were taken on day 1, while the late timepoint samples were taken on days 3 and 4 for wild type and PATS, respectively. Results represent the mean of two technical replicates for each triplicate culture (6 samples total). Error bars represent the standard deviation of triplicate cultures.

As shown in **Figure 4A**, there were no significant differences in chlorophyll or carotenoid content between the PATS and wild-type strains during either sampling timepoint for photoautotrophically grown cultures. This result was somewhat unsurprising considering that cyanobacteria produce chlorophyll and carotenoids on the order of milligrams per gram of dry cell weight, while a cumulative total of only 102 micrograms of patchoulol had been produced by the late sampling timepoint. Similarly, photopigment analysis of mixotrophic cultures did not reveal any significant differences between the wild-type and PATS strains (**Figure 4B**). Between growth conditions, photopigment levels were significantly higher at the late sampling timepoint (p < 0.01) for both wild-type and PATS cultures grown under photoautotrophic conditions compared to cultures grown under mixotrophic conditions (**Figure 4**). Photoautotrophic wild-type cells contained 2.58-fold more chlorophyll and 3.88-fold more carotenoids than wild-type mixotrophic cells (4.13 versus 1.60 mg L^−1^ OD_750_^−1^ chlorophyll (late growth) and 1.32 versus 0.34 mg L^−1^ OD_750_^−1^ carotenoids (late growth)). A difference in photopigment levels was expected because photoautotrophic metabolism relies solely on harvesting light to obtain energy and therefore requires more photopigments to capture light compared to cells grown mixotrophically, which can also obtain energy through glycolysis.

### Bicarbonate Patchoulol Production: High and Low Light

After studying patchoulol production for photoautotrophic and mixotrophic cultures, we were interested to see how light intensity would affect production. Two different light conditions were tested with bicarbonate as the sole carbon source: 20 μE m^−2^ s^−1^ and 100 μE m^−2^ s^−1^. Our initial hypothesis was that higher light would result in higher productivity due to both the higher cell densities achievable and also because the *psbA2* promoter responsible for patchoulol synthase expression is light activated.^4,44^

As expected, the high-light cultures grew to higher cell densities compared to low-light cultures (OD_750_ = 11.11 (wild-type) and 11.76 (PATS) for high-light cultures compared to 8.08 (wild-type) and 5.77 (PATS) for low-light cultures), showing that light was the main factor limiting growth for the low-light condition (**Figure 5 A,C**). Also as expected, cultures grown under high light produced more patchoulol than those grown under low light (**Figure 5**). The final titers of patchoulol were 31.7 μg L^−1^ and 27.6 μg L^− 1^ for cultures grown under high and low light, respectively. It should, however, be noted that low-light cultures were allowed to grow for 12 days (due to slower growth) compared to only 9 days for the high-light cultures. The difference between conditions is more apparent when comparing patchoulol production on day 5 when the high-light cultures entered stationary phase and the production of patchoulol slowed. By day 5, the high-light cultures had produced significantly more patchoulol than the low-light cultures (p < 0.001), 27.31 μg L^−1^ patchoulol versus only 13.45 μg L^−1^ for high and low light cultures, respectively. Despite the higher cumulative production, the specific productivity of the lowlight cultures was greater than the specific productivity of high-light cultures and was maintained over a longer period of time (**Figure 5**). Average specific productivities were 0.60 μg L^−1^ day^−1^ OD_750_^−1^ (over 12 days) and 0.45 μg L^−1^ day^−1^ OD_750_^−1^ (over 9 days) for cultures grown under low and high light, respectively. This explains how the low-light PATS cultures—which reached a maximum OD of only 5.77—produced almost as much patchoulol as the high-light PATS cultures that reached an OD of 11.76. Interestingly, there was no difference in growth between the wild-type and PATS strains when grown under high light (**Figure 5C**), but the PATS strain grew slightly slower than the wild type and reached a lower final OD when grown under low light (**Figure 5A**). One possible explanation could be that the patchoulol synthase diverted flux away from photopigment biosynthesis, which would have had a disproportionally negative effect on low-light cultures compared to high-light cultures (because the low-light cultures were already light-deficient and this was further exacerbated by less efficient biosynthesis of light harvesting pigments due to patchoulol synthase competition for FPP). An alternative reason for the growth inhibition observed could be due to the lack of a functional *psbA2* gene in the PATS strain; however, this seems less likely because the growth inhibition was only observed under low light and not high light, the opposite of what would be expected if that was the cause of the inhibition.

**Figure 5.**
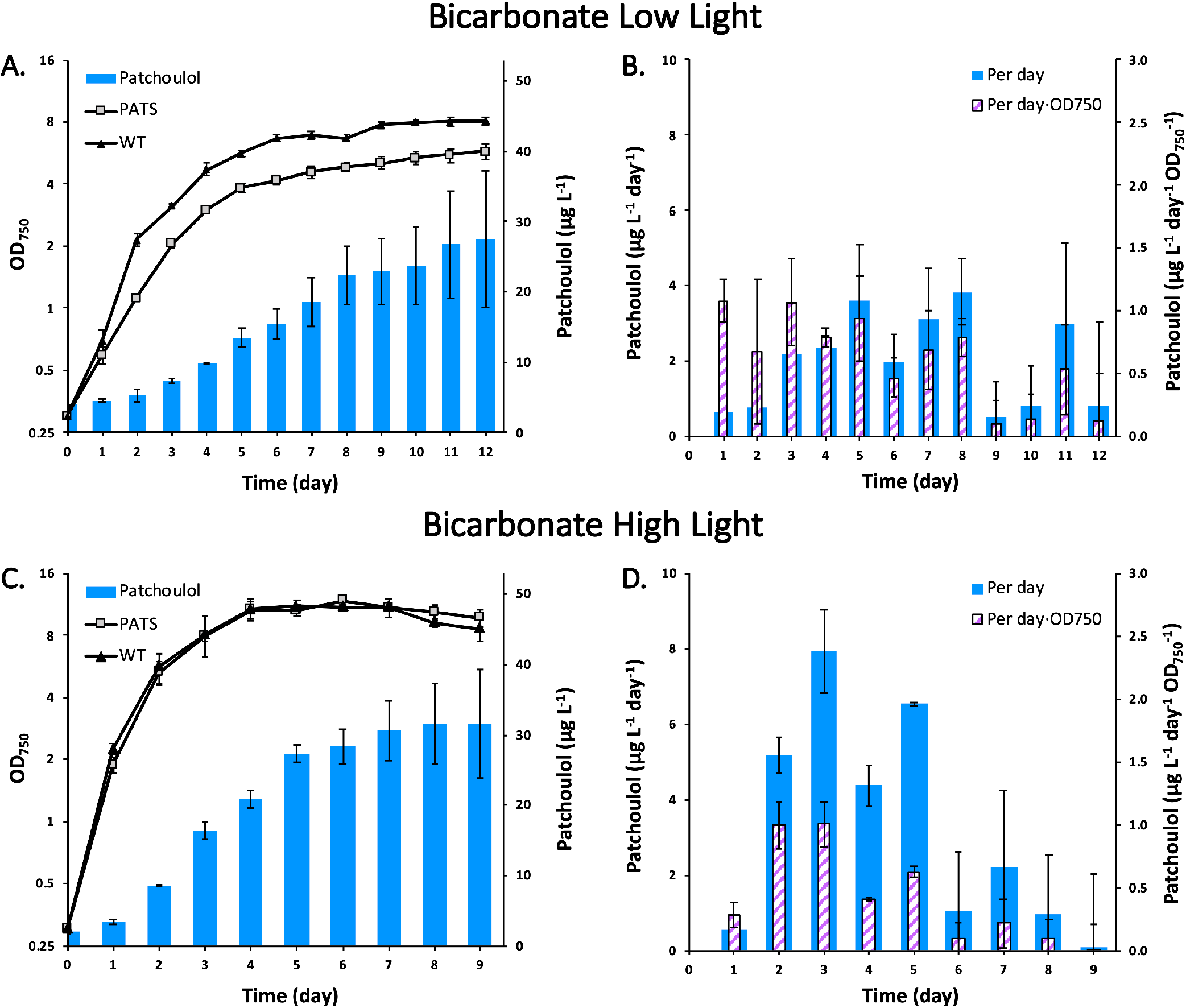
Patchoulol production from *Synechocystis* shake flask cultures with added sodium bicarbonate. (**A**) Growth curves of wild-type and PATS strains under low-light conditions (20 μE m^−2^ s^−1^) and production of patchoulol over time. (**B**) Daily productivity and specific productivity of PATS strain under low light. (**C**) Growth curves of wild-type and PATS strains under high-light conditions (100 μE m^−2^ s^−1^) and production of patchoulol over time. (**D**) Daily productivity and specific productivity of PATS strain under high light. Error bars represent the standard deviation of triplicate cultures. **Note:** growth curves are plotted on a log base 2 scale to better represent exponential growth.

Wild-type cultures grown under low light produced significantly more (p < 0.01) photopigments than high-light cultures at the late growth sampling timepoint (2.02-fold more chlorophyll and 1.45-fold more carotenoids) (**Figure 6**). Interestingly, the PATS strain contained significantly more (p < 0.05) chlorophyll (2.07-fold more) but not significantly more carotenoids for low-light cultures compared to high-light cultures at the late timepoint. The higher photopigment levels for low-light cultures was expected, but the fact that carotenoid levels were almost the same between low-light and high-light PATS cultures at the late sampling timepoint was intriguing. In addition to the PATS strain being slightly growth inhibited under low-light conditions, it had significantly lower levels of both chlorophyll and carotenoids relative to the wild type at both sampling timepoints (**Figure 6A**). Both the growth inhibition and lower photopigment levels relative to the wild type could be explained by patchoulol synthase activity decreasing the intracellular concentration of its substrate, FPP. Since FPP is required for both phytol and carotenoid biosynthesis, this could lead to less efficient photopigment biosynthesis and therefore slower growth under low-light conditions.

**Figure 6.**
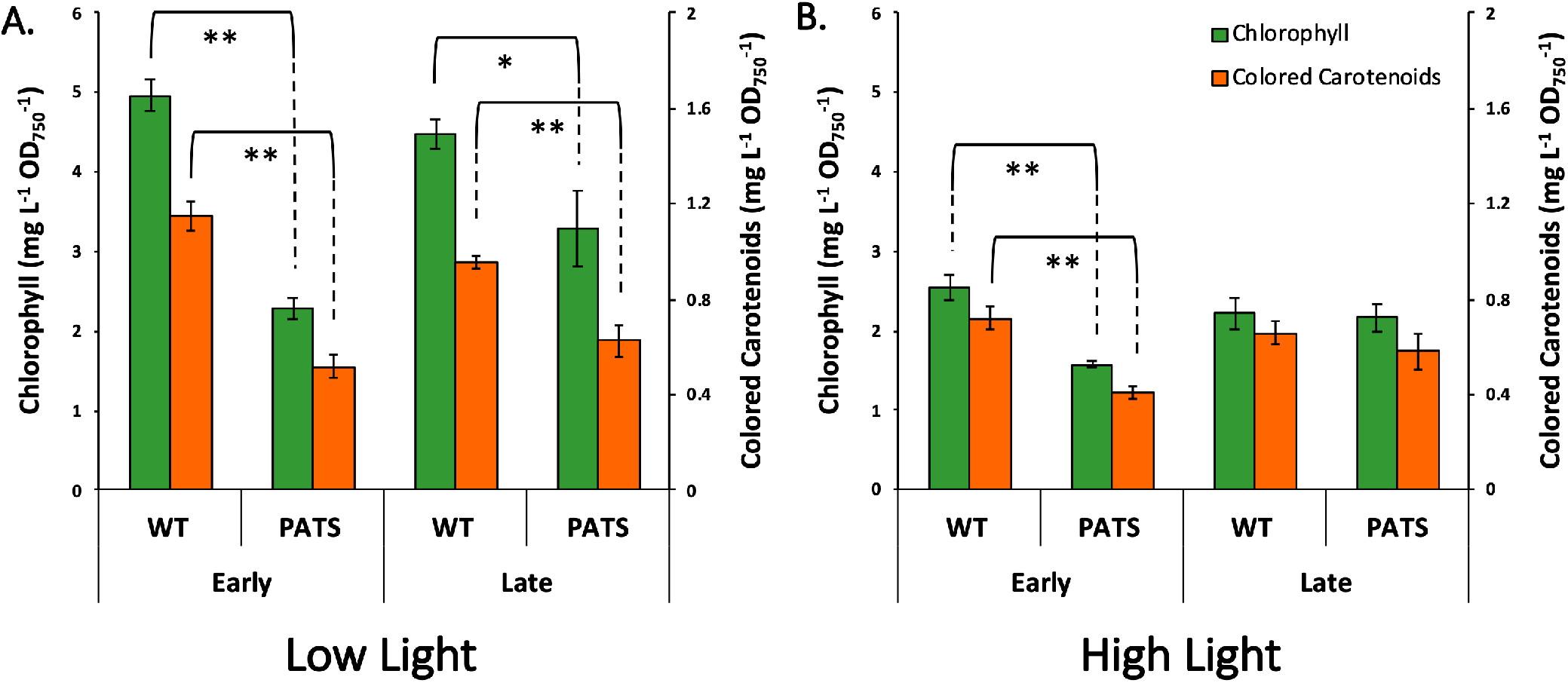
Chlorophyll and colored carotenoid analysis of wild-type and PATS *Synechocystis* strains grown on bicarbonate in shake flasks. (**A**) Cultures grown under low light (20 μE m^−2^ s^−1^). Samples for the early timepoint were taken on day 1 for both the wild type and PATS. Samples taken for the late timepoint were taken on day 6 for the wild type and day 7 for PATS. (**B**) Cultures grown under high light (100 μE m^−2^ s^−1^). Early and late timepoint samples were taken for both strains on days 1 and 4, respectively. Results represent the mean of two technical replicates for each triplicate culture (6 samples total). Error bars represent the standard deviation of triplicate cultures. Asterisks represent statistical significance, where * = p < 0.05 and ** = p < 0.01. Statistical analyses were performed using the average values for each biological replicate (n = 3).

### Understanding the Relationship Between Growth Conditions and Patchoulol Production

Our initial hypotheses were that mixotrophic cultures would produce more patchoulol than photoautotrophic cultures and that high-light cultures would produce more patchoulol than low-light cultures. Our hypotheses were confirmed in one instance and rejected in the other. However, after analyzing the data and comparing the differences in patchoulol and photopigment levels between growth conditions, we noticed that cultures with higher specific patchoulol production also contained more chlorophyll and carotenoids (normalized to OD).

To understand if there was a meaningful relationship between patchoulol and photopigment production that extended across growth conditions, linear regression analysis was performed. Patchoulol was plotted against carotenoids, chlorophyll, and OD (control) for each timepoint that photopigments were quantified (**Figure 7**). The analyses revealed a significant positive correlation between patchoulol production and both chlorophyll and carotenoid production, while the correlation between patchoulol and OD was much weaker. The relationship between patchoulol and carotenoids was slightly stronger than that for patchoulol and chlorophyll (R^2^ = 0.87 and 0.83 for patchoulol versus carotenoids and chlorophyll, respectively). Patchoulol was plotted against OD (**Figure 7C)** as a control to show the relationship between patchoulol production and cell density. This was necessary because some correlation would be expected between patchoulol and most major metabolites simply due to growth of the culture over time and the resulting increase in total biomass. If the strength of the relationship between patchoulol and OD was similar to that of chlorophyll and carotenoids, then photopigment levels would not be useful predictors of patchoulol production. That, however, was not the case, as the coefficient of determination (R^2^ value) of patchoulol with respect to OD was only 0.41, much lower than either value for the photopigments. Furthermore, the p-value obtained from the regression analysis (F-test) of patchoulol versus OD (p = 0.085) did not meet the significance threshold, while p-values were much smaller for the photopigments (p = 0.00074 and 0.0015 for carotenoids and chlorophyll versus patchoulol, respectively). In summary, photopigment production is a much better predictor of patchoulol production than cell density (OD) when comparing between the growth conditions tested in this study. As an additional note, a brief visual inspection of the scatter plot shown in **Figure 7C** may give the impression that there is only one condition (photoautotrophic growth) that weakens the relationship between patchoulol and OD. However, this was shown not to be the case by performing additional regression analyses for each of the three scatter plots with the greatest outlier growth condition removed from each data set. When this was done, the correlations became stronger for each data set as expected, but the correlations between both photopigments and patchoulol were still much stronger (R^2^ = 0.98 and 0.95 for carotenoids and chlorophyll versus patchoulol, respectively) than the correlation between patchoulol and OD (R^2^ = 0.69).

**Figure 7.**
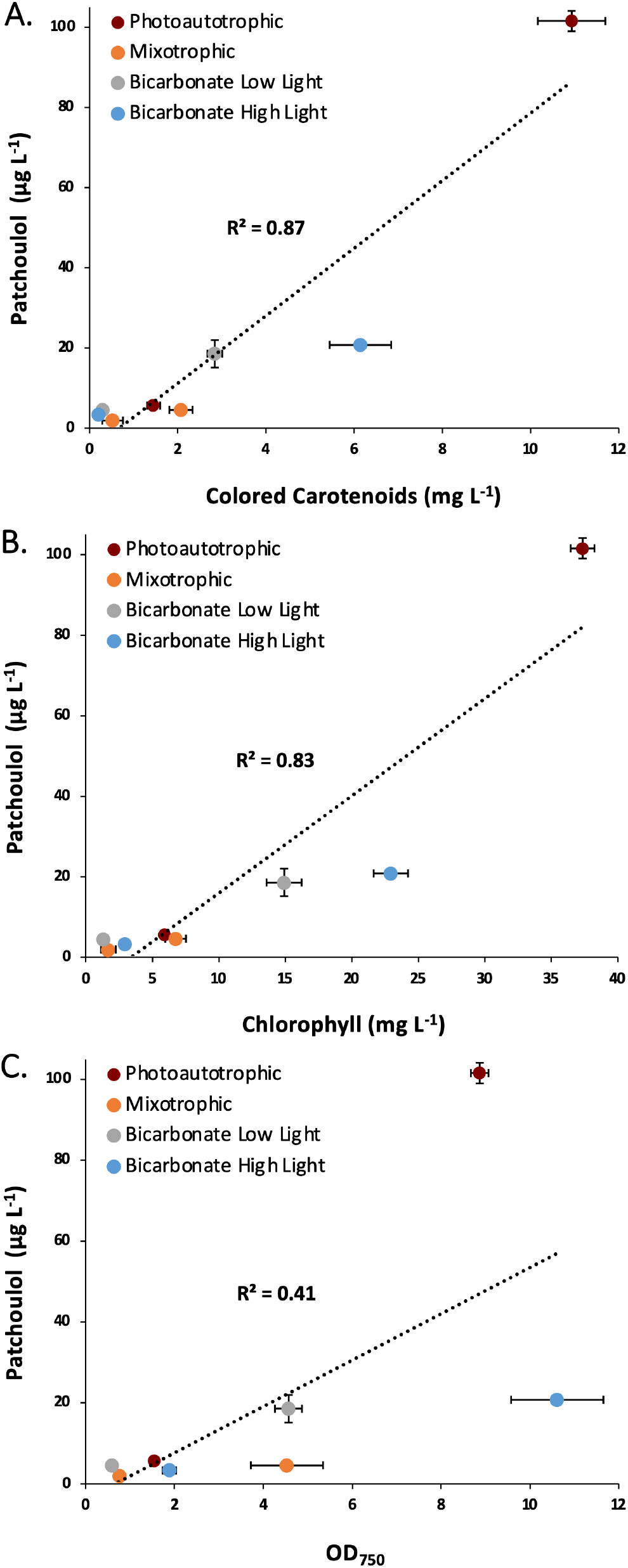
Scatter plots of patchoulol production with respect to colored carotenoids (**A**), chlorophyll (**B**), and OD (**C**). Plots were made using the average values obtained for PATS cultures at each timepoint (early and late) that photopigments were quantified with error bars representing the standard deviation of triplicate cultures. Linear regression analysis was performed using the average value for each condition as a single data point (n = 8).

The relationship between patchoulol production and photopigment production makes sense when considering that carotenoids and the phytol tail of chlorophyll are major terpenoid metabolites in cyanobacteria. Although regulation of the MEP pathway is not fully understood,^13,45^ we hypothesize that flux though the MEP pathway is strongly influenced by growth conditions which promote or suppress the biosynthesis of photopigments that utilize the MEP pathway because they are metabolites that are produced on the order of milligrams per gram of dry cell weight.^4^ If that is the case, then growth conditions which favor photopigment biosynthesis should increase the amount of FPP substrate available for patchoulol synthase (as it is an intermediate in both phytol and carotenoid biosynthesis), increasing patchoulol yields (assuming other variables remain constant).

### Heterologous Terpenoid Production from This Work in Context

The light-dependence results of this study are interesting when compared to a study that expressed a terpene synthase (geranyllinalool synthase) in *Synechocystis* under low and high light (50 and 180 μE m^−2^ s^−1^) and relied on the native MEP pathway for enzyme substrate.^46^ In that study, more than twice as much geranyllinalool was produced under the low-light condition compared to high light. In this work, however, less patchoulol was produced by low-light cultures compared to high-light cultures. There are two differences between studies that may explain this difference. Firstly, the definitions of “low” and “high” light were different between studies. The “low-light” condition in their study consisted of 50 μE m^−2^ s^−1^, which is more than twice the photon flux used for our “low-light” condition. Additionally, the “high-light” condition in their study was 180 μE m^−2^ s^−1^, significantly higher than the “high-light” condition in this study. Their “high-light” condition, which supplied 180 μE m^−2^ s^−1^ light, likely repressed photopigment production even more than what we observed at 100 μE m^−2^ s^−1^, which we propose would then have reduced geranyllinalool yield (based on the correlation we identified in this work between photopigments and terpenoid yield). Secondly, the promoter used in their study was the *cpcB* promoter (the endogenous promoter for the phycocyanin beta subunit), which is downregulated under high-light conditions, while the *psbA2* promoter used in this study is upregulated under high light conditions.^47^ Taken together, the different definitions of low and high light and the differing promoter responses to light likely account for the different terpenoid yields observed. The aforementioned statements are supported by the results of other work where the same promoter and light intensities as this study were used for terpenoid production.^4^ In that study, the high-light condition produced more product (manoyl oxide) than the low-light condition, which was also what we observed. However, there were still some differences between studies. For example, the high-light cultures in their study contained more carotenoids than low-light cultures, the opposite of our findings. Although it can be difficult to directly compare biological studies between laboratories (especially when using different mutant strains), we suggest that these differences may be due to the timing of photopigment quantification during growth, as their study only quantified photopigments at the beginning of growth when all cultures had an OD_750_ < 1.8 at the time of sampling (the lowest being only 0.63).

The patchoulol yield achieved in this study under photoautotrophic conditions (249 μg L^−1^) is comparable to other terpenoids produced in *Synechocystis* strains containing only a heterologous terpene synthase (refer to review articles cited in the introduction). However, the fact that patchoulol synthase has been shown to produce at least 13 sesquiterpenoids other than patchoulol^48^ negatively impacted product yield. Multiple additional sesquiterpenoid products were detected by GC-MS (β-caryophyllene and other unidentified sesquiterpenoids) that were not produced by wild-type cells. This interesting trait of patchoulol synthase may make it well suited for the production of a synthetic essential oil, a concept that has previously been investigated using β-phellandrene synthase enzymes.^49^ Indeed, patchoulol synthase may be particularly suited to this application as in vitro assays have shown that it produces sesquiterpenoids in the same ratios as those found in natural patchouli essential oil.^48^

A recent study reported a yield of 17.3 mg L^−1^ patchoulol from *Synechocystis* using a High-Density Cultivation (HDC) system.^40^ Though an impressive yield, it is based on small-scale (8 mL) cultures grown to an OD_750_ of 32 under 750 μE m^−2^ s^−1^ light, and the authors noted that specific productivity did not increase compared to batch cultures. For small-scale cultures that did not receive increasing light or media replenishment during growth and were supplied 100 μE m^−2^ s^−1^ light (growth conditions comparable to this work), yields of patchoulol were reduced to 590 μg L^−1^, around two-fold greater than the highest yield achieved in this work. Although patchoulol yields were lower in this work, the approach taken was quite different. This work investigated how patchoulol productivity changed as cells used different metabolic growth strategies and made no attempt to optimize conditions to achieve the highest possible yield.

In this work, we showed that there is great variation in terpenoid yields for *Synechocystis* cultures grown under different conditions. In addition to successfully demonstrating the production of the sesquiterpenoid, patchoulol, in *Synechocystis*, we identified a significant positive correlation between photopigment production and heterologous patchoulol production across four different growth conditions. Furthermore, our results demonstrate that photopigment levels are a better predictor of terpenoid yields than cell density when comparing cultures grown under different conditions. Based on these results, we propose a strategy for improving terpenoid production in cyanobacteria through the optimization of growth conditions to produce photopigments, resulting in increased flux through the MEP pathway. Optimized growth conditions can be used alongside other terpenoid production strategies to achieve the highest yields possible.

## METHODS AND MATERIALS

### Strains and Growth Conditions

NEB 5-alpha Competent *E. coli* (High Efficiency) cells (New England Biolabs) were used for plasmid construction. Cultures were grown at 37 °C with shaking at 225 rpm in lysogeny broth (LB) medium (Sigma-Aldrich). The media was autoclaved to sterilize and 1.5% agar (m/v) was added to make plates. Kanamycin (50 μg mL^−1^) was added to the medium when antibiotic selection was required.

A glucose-tolerant wild-type strain of *Synechocystis* sp. PCC 6803 was kindly provided by Dr. Carrie Eckert. General growth of cyanobacterial cultures was performed using BG-11 medium (**Supplemental Table 2**) that was buffered with 10 mM MOPS and brought to a starting pH of 7.6. The media was filter sterilized through a 0.22 μm PES filter. To make plates, 3% agar (m/v) in water (2x concentration) and an equal volume of 2x concentrated BG-11 were autoclaved separately and mixed while still warm before pouring. For general growth of cultures and during transformation experiments, cultures were incubated at 30 °C and irradiated with white light at 20 μE m^−2^ s^−1^. Kanamycin (50 μg mL^−1^) was added to PATS cultures for general growth. A spectrophotometer (Agilent Cary 8454 UV-Visible) was used to measure optical density (OD) at 750 nm to track growth for all cultures.

#### Photoautotrophic and Mixotrophic Growth

Triplicate cultures (70 mL) were grown in BG-11 medium supplemented with 10 mM MOPS buffer (pH = 7.6) in a Multicultivator 1000-OD (Photon Systems Instruments) photobioreactor at 30 °C with 0.4% CO_2_ mixed with air bubbling through cultures and 100 μE m^−2^ s^−1^ LED white light. All cultures had a starting OD of 0.3, and PATS cultures were grown with 50 μg mL^−1^ kanamycin. Photoautotrophic cultures were grown with CO_2_ as the sole carbon source, while mixotrophic cultures had 10 g L^−1^ glucose added to the medium. Samples were taken daily (1 mL) to track glucose consumption for mixotrophic cultures. A biphasic solvent overlay was employed to capture patchoulol by adding 7 mL of filter-sterilized n-dodecane (Acros Organics) to each culture (10% v/v). Optical density of cell cultures was measured daily to track growth, and 100 μL of the dodecane overlay was sampled each day and stored at −80 °C until it was analyzed via GC-MS for patchoulol. At two time points (early and late), 2 mL samples were taken from each triplicate culture for chlorophyll and carotenoid quantification.

#### Bicarbonate Growth

Triplicate cultures (50 mL) were grown in 250 mL unbaffled shake flasks in BG-11 medium supplemented with 50 mM sodium bicarbonate and 30 mM TES buffer with a starting pH of 7.6. Cultures were grown in an incubator at 30 °C with shaking at 100 rpm. Cultures had a starting OD of 0.3, and PATS cultures were supplemented with 50 μg mL^−1^ kanamycin. Cultures were irradiated with LED white light at either 20 μE m^−2^ s^−1^ (low light), or 100 μE m^−2^ s^−1^ (high light). Each day, 2.5 mL of sterile 1 M NaHCO_3_ in 1x BG-11 was added to each triplicate culture to ensure there was a constant supply of bicarbonate. Despite the added buffer, cultures required the pH to be adjusted to ~8 by the addition of concentrated HCl (every 8 hours for the cultures grown under high light and approximately every 12 hours for cultures grown under low light). To capture patchoulol, 5 mL of filter-sterilized n-dodecane (Acros Organics) was added to each culture (10% v/v). OD measurements were taken daily to track cell growth, and 100 μL of the dodecane overlay was sampled each day and stored at −80 °C until it was analyzed by GC-MS at the end of the experiment. At two time points (early and late), 2 mL of each triplicate culture was sampled for chlorophyll and carotenoid quantification.

### Generation of a Patchoulol-producing *Synechocystis* mutant

A patchoulol-producing *Synechocystis* strain was created by knocking out the native *psbA2* open reading frame via double homologous recombination using a suicide vector, pEERM1-PATS, and replacing it with a synthetic, codon optimized patchoulol synthase gene from *Pogostemon cablin* (Genbank Accession no.: AY508730). The patchoulol synthase gene was synthesized and codon optimized for expression in *Synechocystis* by Genewiz (codon optimized DNA sequence given in **Supplementary Information)**.

#### Plasmid Construction

The pEERM1-PATS vector was created by inserting the synthetic patchoulol synthase gene into the pEERM1 plasmid (Addgene plasmid #64024)^4^ using HiFi assembly. First, the pEERM1 backbone was linearized by restriction digestion with *Xba*I. The synthetic patchoulol synthase gene was then PCR amplified using Q5 DNA polymerase with the primers 6803PATS_F and 6803PATS_R (**Supplemental Table 1**) that were designed to have 20 bp 5’ tails that overlapped with the ends of the linearized pEERM1 vector. Both the linearized vector and PCR product were then run on an agarose gel, and the correct sized bands were cut out and purified using a QIAquick Gel Extraction Kit (Qiagen). A HiFi assembly reaction was then performed to join the PCR product with the linear pEERM1 to form the assembled circular plasmid by incubating at 50 °C for one hour using NEBuilder DNA HiFi Assembly Master Mix (New England Biolabs). The HiFi reaction solution (5 μL) was then transformed into 50 μL of NEB 5-alpha Competent *E. coli* (High Efficiency) cells (New England Biolabs) and plated onto LB plates containing 50 μg mL^−1^ kanamycin. After incubating overnight at 37 °C, colonies were screened by colony PCR using the patchoulol synthase HiFi primers aforementioned and were patched out onto fresh LB kanamycin plates. Colonies that were positive following screening were grown up overnight in 10 mL of LB with kanamycin, plasmid DNA was extracted using a QI-Aprep Spin Miniprep Kit (Qiagen), and the purified plasmid DNA was sent to Eurofins Genomics for Sanger sequencing. All plasmids sent for sequencing confirmed correct plasmid assembly without any introduced mutations.

#### Synechocystis Transformation

*Synechocystis* cells were transformed using a modified protocol from Chauvat et al., 1986.^50^ Cells were grown at 30 °C with 20 μE m^−2^ s^−1^ LED white light. Under laminar flow, 10 mL of a culture in exponential growth was harvested, washed with fresh BG-11, and resuspended in 300 μL of BG-11. The cells were divided into 100 μL aliquots and ~5μg of DNA was added for each transformation. The tubes were flicked a few times to gently mix, and the cells were incubated at 30 °C under 50 μE m^−2^ s^−1^ LED white light for 5 hours. Cells were then plated onto BG-11 agar plates and were allowed to recover overnight at 30 °C under 20 μE m^−2^ s^−1^ light. The following day, 1 mL of 50 μg mL^−1^ kanamycin in water was added underneath the agar plate using a pipet to select for successfully transformed cells. Wild-type cells began to die back after a few days, and mutant colonies were visible after about two weeks. Colonies were picked and grown in 5 mL of liquid BG-11 medium with 50 μg mL^−1^ kanamycin at 30 °C with 20 μE m^−2^ s^−1^ light. To screen colonies that grew in liquid culture, 100 μL samples were taken and centrifuged briefly to pellet the cells. The supernatant was removed with a pipet and the cells were resuspended in 100 μL of water and incubated at 99 °C for 10 minutes to lyse cells. The tubes were centrifuged to remove cellular debris, and 1 μL of the cell boilate was used as DNA template for PCR to screen for successful transformants using the primers 6803psbA2_F and 6803psbA2_R (initial screening PCR and primer sequences are provided in **Supplemental Figure 2 and Supplemental Table 1**, respectively). Primers annealed outside of the inserted region in the *Synechocystis* genome, producing a smaller (1721 bp) amplicon for wild-type cells and a larger amplicon (3480 bp) for the PATS strain.

### Patchoulol Quantification

Samples (100 μL) of the dodecane overlay were taken from each culture daily and stored at −80 °C until all samples were collected. The concentration of patchoulol in the dodecane layer was measured by GC-MS and calculated using a standard curve (R^2^ > 0.99) made using a patchoulol primary reference standard (Sigma-Aldrich) in dodecane. The volumes removed due to sampling from dodecane overlays and cell cultures were accounted for during patchoulol quantification. At the end of each experiment, the culture and dodecane were poured into a graduated cylinder to measure the remaining volume and correct for evaporation. The corrected values were used for calculating metabolite concentrations, though the difference was almost negligible.

Patchoulol in dodecane was quantified via GC-MS (Agilent 7890A gas chromatograph interfaced with an Agilent 5975C inert MSD with Triple-Axis Detector). Samples (1 μL) were injected in splitless mode at an injector temperature of 220 °C and were separated using an Agilent HP-5MS column (30 m length, 0.250 mm internal diameter, 0.25 μm film thickness). The initial oven temperature was 100 °C for 3 min and was ramped up to 200 °C at a rate of 9 °C/min before ramping up to 300 °C at a rate of 100 °C/min, where it was held for 5 minutes for a total run time of 20.11 minutes. Helium (99.9999 %) was used as the carrier gas (1.0 mL/min. constant flow). The ion source temperature was 220 °C. The mass spectrometer was operated at 70 eV in the electron ionization mode and the mass range scanned was 40–400 m/z. Patchoulol was identified by comparison of retention time and mass spectra with a primary reference standard (Sigma-Aldrich) and was quantified using a standard curve (R^2^ > 0.99).

### Photopigment Quantification

Chlorophyll and colored carotenoids were quantified based on previously established methods.^4,51^ Briefly, two 1 mL samples were taken from each triplicate culture during growth and were placed in 1.5 mL microcentrifuge tubes. The samples were pelleted by centrifugation at 16900 g for 5 minutes, the supernatant was discarded, and the cells were placed in a −20 °C freezer overnight. The cells were allowed to thaw and were vortexed to loosen the pellet. They were then resuspended in 1 mL of N-N-Dimethlyformamide (DMF) (≥ 99%, Sigma-Aldrich) and were incubated at room temperature in the dark for 5 minutes. The cells were pelleted again by centrifugation at 16900 g for 5 minutes. The supernatant was placed in quartz cuvettes, and the absorbance was measured at 461 nm and 664 nm using a spectrophotometer (Agilent Cary 8454 UV-Visible). Concentrations of chlorophyll and colored carotenoids were calculated using the following equations: chlorophyll [μg/mL] = OD_664_ × 11.92; colored carotenoids [μg/mL] = (OD_461_ – (OD_664_ × 0.046)) × 4. Statistical significance was determined using a two-tailed t-test (assuming unequal variances).

### Residual Glucose Quantification

Residual glucose was quantified from 1 mL samples taken from mixotrophic cultures that had been centrifuged at 16900 g for 5 minutes to obtain the supernatant. Supernatant samples were filtered through a 0.22 μm filter and stored at −20 °C until all samples had been collected. Samples were thawed, filtered again through a 0.22 μm filter, and loaded onto a HPLC (Agilent 1260 Infinity II LC) equipped with a refractive index detector (RID). Samples were analyzed by isocratic elution using a 5 mM H_2_SO_4_ mobile phase at a constant flow rate of 0.5 mL min^−1^ on a REZEX ROA-organic acid H^+^ (8%) LC column (300 mm × 7.8 mm, Phenomenex). The column temperature was maintained at 55 °C and the refractive index detector was operated at 35 °C. Glucose concentration was calculated using a standard curve (R^2^ > 0.99).

## Supporting information

S.I. Synechocystis Terpenoid Production Different Growth Conditions

## ASSOCIATED CONTENT

### Supporting Information

Supplementary information accompanies this paper and is available to download.

### Author Contributions

R.H. conceived the idea for the project, performed the experiments, and wrote the manuscript. S.B. helped guide the project and edit the manuscript.

### Funding Sources

This work was supported by the Green Chemicals Beacon, University of Nottingham. R.H. was supported by a Fulbright-University of Nottingham Postgraduate Award.

## ACKNOWLEDGMENT

We would like to thank the Green Chemicals Beacon for funding this work. We would also like to thank Dr. Stephen Hall for his assistance with GC-MS method development and Dr. Andrew Yiakoumetti for his assistance with glucose quantification by HPLC. R.H. is grateful to the US-UK Fulbright Commission and the University of Nottingham for funding.

## ABBREVIATIONS

PATS: patchoulol-producing *Synechocystis* strain
FPP: farnesyl diphosphate
G3P: glyceraldehyde-3-phosphate
MEP pathway: methylerythritol phosphate pathway
OD: optical density
GCMS: gas chromatography-mass spectrometry
HPLC: high performance liquid chromatography

