## Supplementary material for "Engineered Terpenoid Production in *Synechocystis* sp. PCC 6803 Under Different Growth Conditions": S.I. Synechocystis Terpenoid Production Different Growth Conditions

**Supplementary Information**

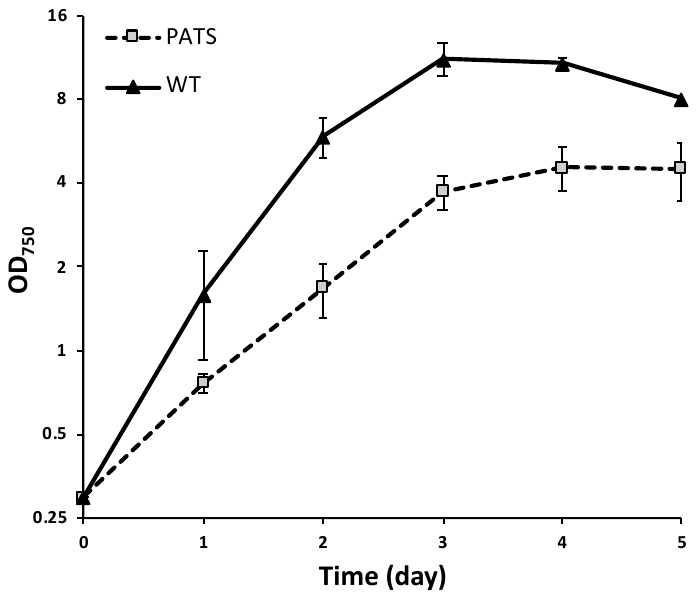

**Supplemental Figure 1** Preliminary experiment showing a growth inhibition for the PATS strain relative to the wild type under mixotrophic conditions. Growth curves were plotted on a log base 2 scale to better represent exponential growth.

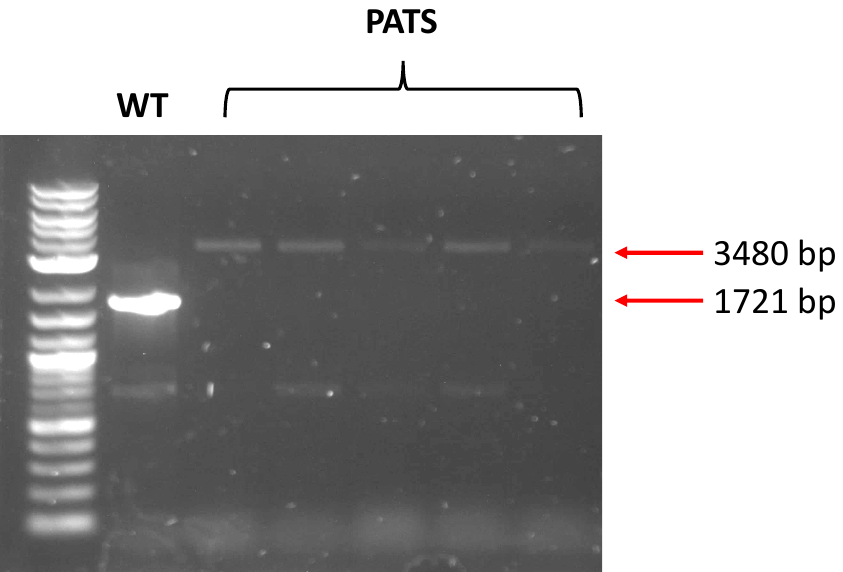

**Supplemental Figure 2** Agarose gel showing initial screening of fully segregated Synechocystis PATS colonies that grew in liquid medium compared to the wild type (using primers 6803psbA2_F and 6803psbA2_R). DNA was amplified from cell boilate. PCR from the PATS and wild-type strains gave bands of the expected sizes, 3480 and 1721 bp, respectively. No wild-type amplicon was detected for any of the PATS isolates.

**Supplemental Table 1** Primer sequences used for construct assembly and PCR screening

| **Primer Name** | **Sequence (5’**$\boldsymbol{\to}$**3’)** |
| --- | --- |
| 6803PATS_F | CAAATACATTAGTGGAGGTTATGGAATTATATGCCCAATC |
| 6803PATS_R | tgcagcggccgctactagttTTAATAAGGCACGGGATG |
| 6803psbA2_F | CAGTTGTGTCGCCCCTCTAC |
| 6803psbA2_R | CAAATCTAACGAACCGCCCC |

**Supplemental Table 2** Composition of BG-11 Medium

| **Component** | **Amount (per Liter)** |
| --- | --- |
| NaNO_3_ | 1.5 g |
| MgSO_4_⋅7H_2_O | 0.075 g |
| CaCl_2_⋅2H_2_O | 0.036 g |
| K_2_HPO_4_⋅3H_2_O | 0.04 g |
| **Trace Elements (1000x)** | 1 mL |
| **Trace Elements (1000x)** | **Amount (per Liter)** |
| Citric Acid | 6 g |
| Ferric Ammonium Citrate | 6 g |
| H_3_BO_3_ | 2.86 g |
| MnCl_2_⋅4H_2_O | 1.81 g |
| Na_2_EDTA·2H_2_O | 1 g |
| NaMoO_4_⋅2H_2_O | 0.391 g |
| ZnSO_4_⋅7H_2_O | 0.222 g |
| CuSO_4_⋅5H_2_O | 0.079 g |
| Co(NO_3_)_2_⋅6H_2_O | 0.049 g |

**Note:** pH was adjusted to 7.6 prior to filter sterilization

**Patchoulol synthase codon optimized DNA sequence**

ATGGAATTATATGCCCAATCCGTGGGTGTGGGTGCTGCTAGTCGCCCCTTAGCCAATTTTCATCCTTGTGTGTGGGGTGACAAGTTCATCGTGTACAACCCCCAGAGTTGTCAAGCCGGCGAGCGGGAGGAAGCCGAGGAATTAAAGGTGGAGTTGAAACGGGAGTTGAAAGAGGCCTCCGATAACTACATGCGCCAGTTGAAGATGGTGGACGCCATCCAGCGCTTGGGCATCGACTATTTGTTCGTTGAGGACGTGGACGAGGCCTTGAAAAATTTGTTCGAGATGTTCGATGCCTTTTGTAAGAATAATCACGACATGCACGCCACTGCCTTGTCCTTTCGGTTGTTGCGGCAACATGGTTACCGCGTTAGTTGTGAGGTGTTTGAGAAATTCAAGGACGGCAAGGACGGCTTCAAAGTTCCCAATGAGGACGGCGCTGTGGCTGTGTTAGAGTTCTTTGAGGCCACCCATTTGCGGGTTCATGGTGAGGATGTGTTGGACAACGCCTTCGACTTCACCCGGAACTACTTAGAGTCCGTGTATGCCACCTTGAACGATCCTACCGCCAAGCAAGTGCACAACGCCTTGAACGAATTCAGTTTTCGCCGCGGTTTACCCCGGGTGGAGGCCCGGAAATACATTAGTATTTATGAACAGTACGCCTCCCACCACAAGGGCTTATTGAAATTGGCTAAGTTGGACTTCAACTTGGTGCAAGCCTTGCATCGCCGGGAATTGAGTGAGGATTCCCGCTGGTGGAAGACCTTGCAAGTGCCCACCAAGTTGAGTTTCGTGCGGGACCGGTTAGTGGAGAGTTATTTCTGGGCCTCCGGCTCCTACTTCGAGCCCAACTATTCCGTGGCCCGGATGATCTTGGCCAAGGGTTTGGCCGTGTTATCCTTGATGGACGACGTGTATGACGCCTATGGCACCTTCGAAGAATTGCAGATGTTCACCGATGCTATTGAACGCTGGGATGCCAGTTGTTTAGATAAGTTGCCCGACTATATGAAGATCGTTTACAAGGCCTTATTAGATGTGTTTGAGGAAGTGGATGAGGAGTTGATTAAATTGGGCGCCCCCTACCGGGCCTATTACGGCAAGGAAGCCATGAAGTACGCCGCTCGGGCTTATATGGAAGAAGCCCAATGGCGGGAGCAGAAGCATAAACCCACCACCAAGGAGTATATGAAATTAGCTACTAAGACTTGTGGCTATATTACTTTGATTATCTTGAGTTGTTTGGGCGTGGAAGAGGGCATCGTGACCAAGGAGGCCTTCGATTGGGTTTTCTCCCGGCCCCCCTTCATCGAGGCCACCTTGATTATCGCCCGCTTGGTGAACGATATTACCGGCCACGAGTTCGAAAAGAAACGGGAACACGTGCGCACCGCCGTGGAATGCTACATGGAGGAACACAAGGTGGGCAAACAAGAAGTGGTTTCCGAGTTCTACAACCAGATGGAGTCCGCTTGGAAGGACATTAACGAGGGTTTCTTGCGGCCCGTGGAATTTCCCATTCCCTTGTTGTACTTGATTTTAAACTCCGTGCGGACCTTGGAGGTGATCTATAAGGAGGGCGATTCCTACACCCATGTGGGCCCCGCCATGCAGAACATTATCAAACAATTGTATTTACATCCCGTGCCTTATTAA
